# RepB C-terminus mutation of an ori pRi vector affects plasmid copy number in *Agrobacterium* and transgene copy number in plants

**DOI:** 10.1101/365320

**Authors:** Zarir Vaghchhipawala, Sharon Radke, Ervin Nagy, Mary L. Russell, Susan Johnson, Stanton B. Gelvin, Larry Gilbertson, Xudong Ye

## Abstract

A native *repABC* replication origin, ori pRi, was previously reported as a single copy plasmid in *Agrobacterium tumefaciens* and can improve the production of transgenic plants with a single copy insertion of transgenes when it is used in binary vectors for *Agrobacterium-mediated* transformation. A high copy ori pRi variant plasmid, pTF::Ri, which does not improve the frequency of single copy transgenic plants, has been reported in the literature. Sequencing the high copy pTF::Ri *repABC* operon revealed the presence of two mutations: one silent mutation and one missense mutation that changes a tyrosine to a histidine (Y299H) in a highly conserved area of the C-terminus of the RepB protein (RepB^Y299H^). Reproducing these mutations in the wild-type oriRi binary vector showed that *Agrobacterium* cells with the RepB^Y299H^ mutation grow faster on both solidified and in liquid medium, and have higher plasmid copy number as determined by ddPCR. In order to investigate the impact of the RepB^Y299H^ mutation on transformation and quality plant production, the RepB^Y299H^ mutated ori pRi binary vector was compared with the original wild-type ori pRi binary vector and a multi-copy oriV binary vector in canola transformation. Molecular analyses of the canola transgenic plants demonstrated that the multi-copy ori pRi with the RepB^Y299H^ mutation in *Agrobacterium* cells lost the advantage of generating high frequency single copy, backbone-free transgenic plants compared to using the single copy wild-type ori pRi binary vector.

## Introduction

Single copy, vector backbone-free transgene events are required by government regulatory agencies for commercial transgenic product deployment, and are preferred for many other biotechnology applications. This is because most vector backbones contain antibiotic genes and other not fully characterized sequences, and that multi-copy transgenes may have reduced or silenced gene expression [1–3]. Efforts have been made to increase single copy transgene frequency, and to eliminate the presence of vector backbone sequences in transgenic plants using particle bombardment or *Agrobacterium-mediated* transformation [4–9]. Lowering DNA loading in particle bombardment [4–8], or using a single copy binary vector rather than multi-copy vectors in *Agrobacterium* transformation [9]. can significantly increase single copy transgenic plant production.

*Agrobacterium*-mediated plant transformation often uses a binary vector to maintain the plasmid in *Escherichia coli* for gene cloning manipulation and in *Agrobacterium* cells to deliver the T-strand into plant cells. Different origins of replication control the binary vector copy number in *Agrobacterium* which can further affect T-strand processing and integration into the plant genome. The broad host range plasmid RK2-derived origin of replication (RK2 oriV, incPα group, GenBank accession #J01780) replicates to approximately 10 copies/Agrobacterium chromosome [10], whereas the ori pRi (oriRi, GenBank accession # X04833), a plasmid with the origin of replication from the *Agrobacterium rhizogenes* plasmid pRiA4b, replicates the plasmid as 1 copy/Agrobacterium chromosome [11–13]. Native oriRi-based binary vectors produce a higher percentage of single copy, backbone free transgenic plants across multiple crops than do multi-copy binary vectors [9]. A new variant of the ori pRi replicon, named pTF::Ri, has been GGCAAGAACAATCCTCAAACCAreported at an estimated 15-20 copies of binary plasmid per *Agrobacterium* cell. This ori provided no advantage in single copy transgenic plant production [14].

The oriRi plasmid contains the *repABC* operon encoding the RepA and RepB proteins for plasmid partitioning, and the replication initiator protein RepC for DNA synthesis [15,16]. A small RNA, which binds to the region of *repABC* mRNA between *repB* and *repC*, tightly regulates *repC* expression to control plasmid copy number and determines plasmid incompatibility [17,18]. RepB proteins bind specifically to *parS*-like sequences, and are required for proper segregation of replicated plasmids to daughter cells. RepB can form dimers and oligomers in solution. The middle part of RepB is responsible for *parS* binding and the C-terminus determines RepB dimerization/oligomerization [19]. The *parS* binding site identified in the plasmid pRiA4b is 28 bp downstream from the end of the *repC* ORF [20].

To address the discrepancy between the oriRi [9] and pTF::Ri [14] results, we sequenced the high copy plasmid pTF::Ri. Sequence analysis of the entire plasmid resulted in the identification of two point mutations in the *repB* gene of the oriRi plasmid. In this manuscript, we characterize these two oriRi *repB* mutations for their effect on plasmid copy number in *Agrobacterium*, and their impact on the frequency of single copy, backbone-free transgenic canola plant production.

## Materials and Methods

### Plasmid sequence and construction

The high copy oriRi plasmid pTF::Ri [14] was sequenced and assembled by the Monsanto sequence center. This new sequence was aligned with the published pRiA4b sequence (GenBank accession # X04833) using the EMBOSS::Needle program (http://www.ebi.ac.uk/Tools/emboss/align/index.html).

The oriRi plasmid pMON83937 [9] was used for all site-directed mutagenesis experiments using the QuickChange II protocol (Agilent Cat # 200521). The primers 5’ GATCATGTGCCAGCGCTG**<u>cAT</u>**CAAGCGTACCACGCTG 3’ (forward) and 5’ CAGCGTGGTACGCTTG**ATg**CAGCGCTGGCACATGATC 3’ (reverse) were used to generate the RepB^Y299H^ mutation, whereas the primers 5’ GCAGTTTTCTCGAGAG**<u>ATc</u>**GTCATCGCCGCGATGTCG 3’ (forward) and 5’ CGACATCGCGGCGATGAC**gAT**CTCTCGAGAAAACTGC 3’ (reverse) were used to produce the *repB* T486C silent mutation. Ten ng of pMON83937 was used in a 50 μl reaction volume and amplified for 25 cycles with 5 min elongation time. The reaction mixture was incubated with *Dpn*I to remove the template and transfected into *E. coli* DH10B competent cells to recover the plasmids. The resultant plasmids were extracted using a Qiagen maxiprep kit and verified by full plasmid sequencing.

### *Agrobacterium* DNA extraction and plasmid copy number determination

*Agrobacterium* total DNA was prepared according to Estrella et al. [21] with modifications. Briefly, *Agrobacterium* containing the corresponding plasmids was grown in LB medium at 30°C with 50 mg/L spectinomycin selection. When the A_600_ reached approximately 1.0, 1 ml of *Agrobacterium* cells was harvested into an Eppendorf tube and centrifuged at 15000 rpm for 5 min. The pellet was suspended in 300 μl P1 buffer (from Qiagen Maxiprep kit with RNaseA), 100 μl N-lauroylsarcosine (5% solution in TE) and mixed well, followed by addition of 100 μl Pronase (Calbiochem, 2.5 mg/ml solution in TE buffer). The solution was incubated at 37°C for ~2 hours. The lysate was sheared by passing through a 1 ml pipet tip ~10 times, extracted twice with an equal volume of phenol/chloroform (pH 8), and then twice with an equal volume of chloroform. Total DNA from the aqueous phase was precipitated by addition of an equal volume of isopropanol at room temperature, followed by centrifugation at 15000 rpm at 4°C for 10 min. The DNA pellet was washed with 70% ethanol, air-dried, suspended in 100 μl water, and quantified by a Thermo Fisher Nanodrop instrument.

Plasmid copy number was determined using droplet digital PCR (ddPCR) according to Jahn et al. [22] with minor modifications. Primers and MGB (minor grove binding) probes were designed using Primer Express 3.0 (Thermo Fisher Scientific, www.thermofisher.com). The plasmid-specific Taqman assay was designed to target a β-glucuronidase (GUS) expression cassette included in all plasmids. The native *Agrobacterium* gene *lipA* (GenBank AE007869.2) was used as a chromosomal reference template. The *gusA* and *lipA* probes were labeled with FAM and VIC fluorophores, respectively. The primers 5’ AGCCTTCCACGCCTTTCCT 3’ (forward) and 5’ CCGCTTTTCCCGACGAT 3’ (reverse) were used to amplify a *lipA* short sequence, and the MGB (minor grove binding) Taqman probe VIC-5’ CCACCTTGCAGATTG 3’-MGB was used for real time *lipA* amplification. The primers 5’ ACCGAATACGGCGTGGATAC 3’ (forward) and 5’ TCCAGCCATGCACACTGATAC 3’ (reverse) were used to amplify a *gusA* short sequence, and the MGB Taqman probe 6FAM-5’ TGTACACCGACATGTGG 3’-MGB was used for real time *gusA* amplification. Duplex PCR reactions were performed following the manufacturer’s recommendations [23]. The 20 μl reaction volume included 0.24 pg/μl total template DNA. The concentrations of each primer and probe were 0.9 μM and 0.25 μM, respectively. Following an initial denaturation step at 95°C, a two-step amplification cycle (94°C and 59°C) was repeated 40 times. Four technical replicates were taken for each DNA sample. Template concentrations were calculated by the Quantasoft 1.4 software (Bio-Rad, www.bio-rad.com) and were given as copy/μl values. Plasmid copy numbers were calculated by normalizing the *gusA* template concentrations to the *lipA* template concentrations.

### Canola transformation

The pMON83937 RepB^Y299H^ mutation plasmid (designated as pM0N138207), the original oriRi binary vector pMON83937, and the oriV binary vector pMON67438 [9] were transferred into *Agrobacterium tumefaciens* ABI and plasmid-containing bacteria were selected using 50 mg/L spectinomycin and 50 mg/L kanamycin. Canola (*Brassica napus* L, cultivar Ebony) transformation, CP4 transgene copy number determination, and statistical analysis have been described previously [9].

### Statistical analysis

All experiments were carried out under identical conditions and repeated at least three times. Canola explants were randomized before *Agrobacterium* inoculation. Data are presented as averages of the mean of three independent experiments with standard deviation. Significant differences among the results were assessed by Analysis of Variance (ANOVA).

## Results and Discussion

### pTF::Ri RepB protein sequence has a mutation in a highly conserved residue at the C-terminus

Sequence alignment of the sequence of pTF::Ri with the GenBank ori pRi (# X04833) indicates that there are two T to C point mutations in the *repB* gene of the oriRi: one a T to C silent mutation at 486 bp of the *repB* coding region (*repB* T486C), and the other a T to C point mutation at 895 bp of the *repB* gene leading to a Y299H mutation (RepB^Y299H^; see the *repB* alignment in **Supporting information**). The RepB^Y299H^ mutation results in a change from a hydrophobic to a hydrophilic amino acid in the C-terminal region (Figure 1).

**Figure 1:**
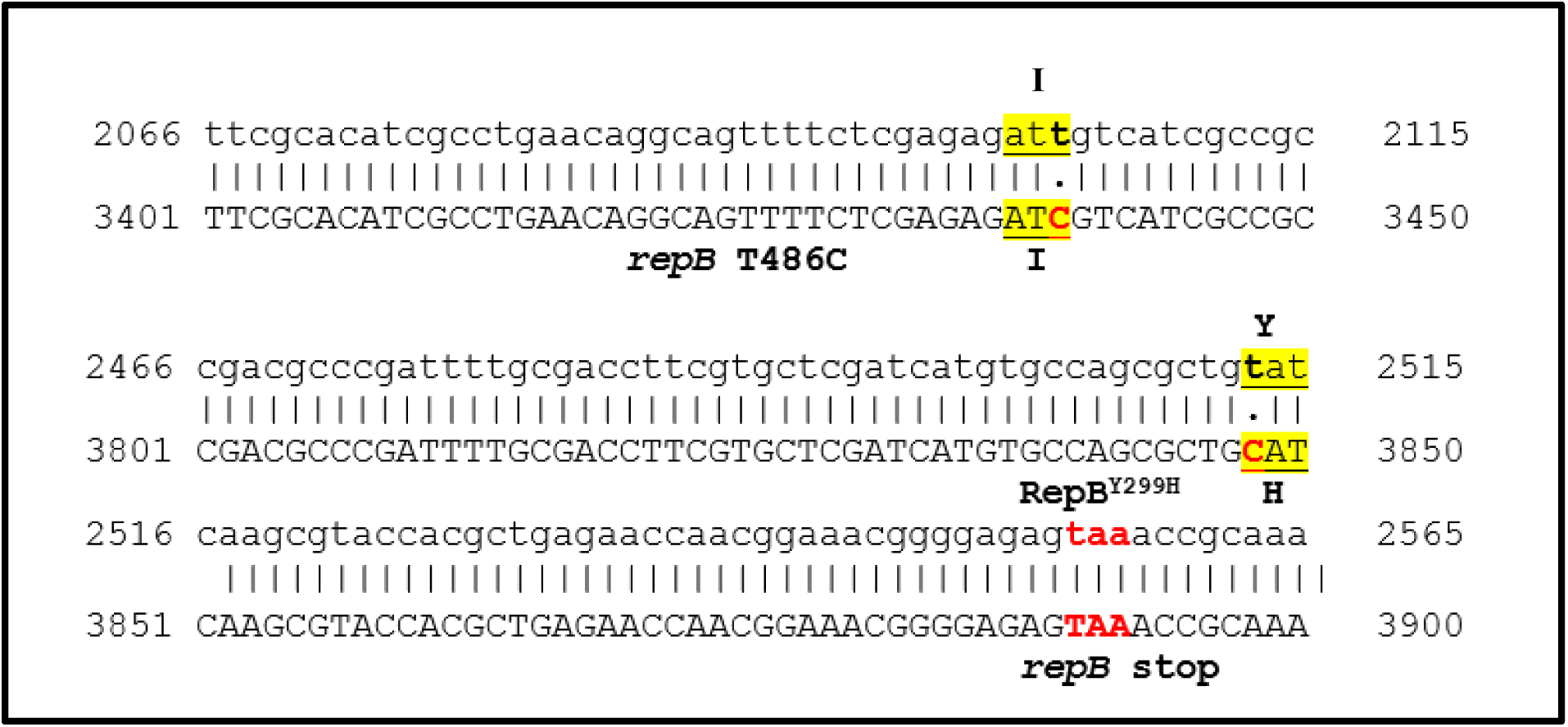
Sequence alignment between part of the genes encoding wild-type *repB* and *repB* from the pTF::Ri binary vector. The T to C changes are highlighted. The *repB* stop codon is marked in red. Upper strand: wild-type *repB* from GenBank X04833; lower strand: pTF::Ri sequence.

To investigate the potential role of the RepB C-terminal sequence, we aligned 26 RepB C-termini from GenBank (Figure 2). The oriRi RepB Y299 is conserved in the majority of *repABC* replicons, indicating that it may play an important role in RepB function.

**Figure 2:**
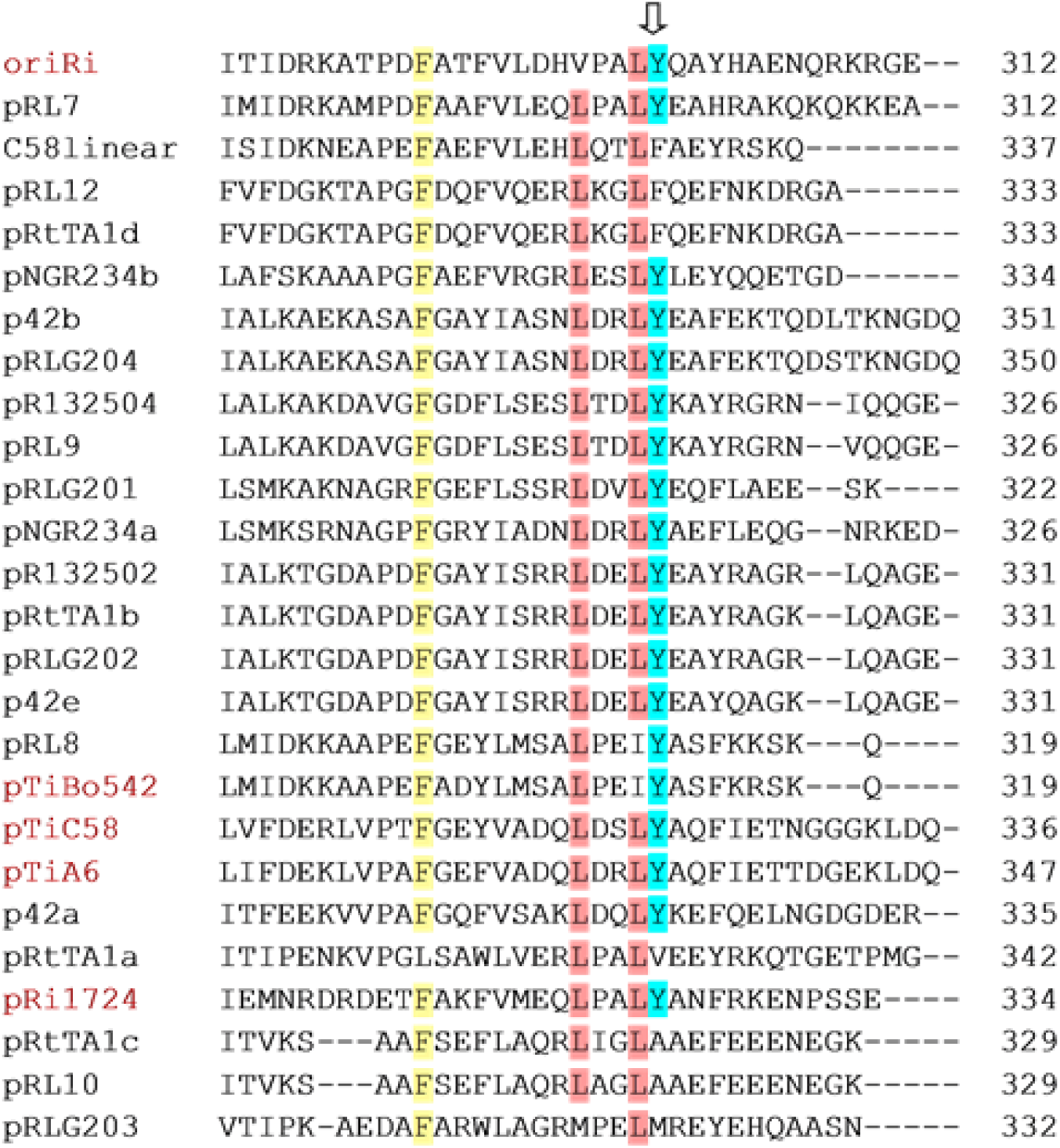
Alignment of the oriRi RepB C-terminus with other RepB C-termini. Arrow indicates the Y299H point mutation. The number at right indicates the last amino acid of each RepB protein. (GenBank accession: oriRi, # X04833; pRL7, *Rhizobium leguminosarum* bv. *viciae* plasmid #AM236081; C58linear, *Agrobacterium tumefaciens* str. C58 linear chromosome # AE007870.2; pRL12, *Rhizobium leguminosarum bv. viciae* plasmid pRL12 # AM236086; pRtTA1d, *Rhizobium leguminosarum bv. trifolii* strain TA1 plasmid # FJ592234; pNGR234b, *Sinorhizobium fredii* NGR234 plasmid pNGR234b # CP000874; p42b, *Rhizobium etli* CFN 42 plasmid p42b # NC_007763; pRLG204, *Rhizobium leguminosarum bv. trifolii* WSM2304 plasmid pRLG204 # NC_011371; pR132504, *Rhizobium leguminosarum bv. trifolii* WSM1325 plasmid pR132504; pRL9, *Rhizobium leguminosarum bv. viciae* plasmid pRL9 # NC_008379; pRLG201, *Rhizobium leguminosarum bv. trifolii* WSM2304 plasmid pRLG201; pNGR234a; *Sinorhizobium fredii* NGR234 plasmid pNGR234a # U00090; pR132502, *Rhizobium leguminosarum bv. trifolii* WSM1325 plasmid pR132502; pRtTA1b, *Rhizobium leguminosarum bv. trifolii* strain TA1 plasmid pRtTA1b # FJ592235; pRLG202, *Rhizobium leguminosarum bv. trifolii* WSM2304 plasmid pRLG202 3 # NC_011366; p42e, *Rhizobium etli* CFN 42 plasmid p42e # NC_007765; pRL8, *Rhizobium leguminosarum bv. viciae* plasmid pRL8 # NC_008383; pTiBo542, *Agrobacterium tumefaciens* Ti plasmid pTiBo542 # DQ058764; pTiC58, *Agrobacterium tumefaciens* str. C58 Ti plasmid # NC_003065; pTiA6, *Agrobacterium tumefaciens* octopine-type Ti plasmid # AF242881; p42a, *Rhizobium etli* CFN 42 plasmid p42a # CP000134; pRtTA1a, *Rhizobium leguminosarum bv. trifolii* TA1 plasmid pRtTA1a # HM032068; pRi1724, *Agrobacterium rhizogenes* plasmid pRi1724 # NC_002575; pRtTA1c, *Rhizobium leguminosarum bv. trifolii* TA1 plasmid pRtTA1c # EU555187; pRL10, *Rhizobium leguminosarum bv. viciae* plasmid pRL10 # NC_008381; pRLG203, *Rhizobium leguminosarum bv. trifolii* WSM2304 plasmid pRLG203 # NC_011370). Yellow, red, and blue shading indicate generally conserved amino acids.

### The RepB^Y299H^ mutation enhances culture growth and increases plasmid copy number in *Agrobacterium*

The *repABC* sequence in the oriRi replicon of pMON83937 used in a prior study [9] is identical to that of the oriRi GenBank reference sequence. To address the question of whether the presence of either or both mutations leads to the increase in plasmid copy number in *Agrobacterium* [14], we carried out site-directed mutagenesis to recapitulate the T to C base changes at 486 bp and 895 bp in pMON83937. These two plasmids were designated pMON83937-*repB*^T486C^ and pMON83937-RepB^Y299H^, respectively. Both plasmids and the control pMON83937 were separately introduced into the nopaline-type strain *Agrobacterium tumefaciens* ABI [9] by electroporation, and the cultures were spread on LB medium containing 50 mg/L spectinomycin and 10 mg/L gentamicin. After 72 hours, colonies harboring RepB^Y299H^ were much larger than those containing the control plasmid or the *repB* T486C silent mutation (Figure 3). The faster growth could result from elevated plasmid copy number, providing a growth advantage from the multiple copies of the *aadA* gene [24]. To investigate this phenomenon further, at least two individual colonies from two sequence confirmed plasmids per mutation were suspended into 5 ml LB medium with the requisite antibiotics at A_660_=0.5, an equal volume of culture was inoculated into 50 ml medium, and the ODs recorded after overnight growth (21 hrs). Over three repetitions of this experiment, the cultures with the RepB^Y299H^ mutation consistently showed a 120-125% higher A_660_ over the control, whereas the A_660_ of the *repB*^T486C^ cultures showed 90% growth compared to the control (Figure 3B). Cultures containing the RepB^Y299H^ mutation reached A_660_=1.0 about 3 hours earlier than did the control culture.

**Figure 3:**
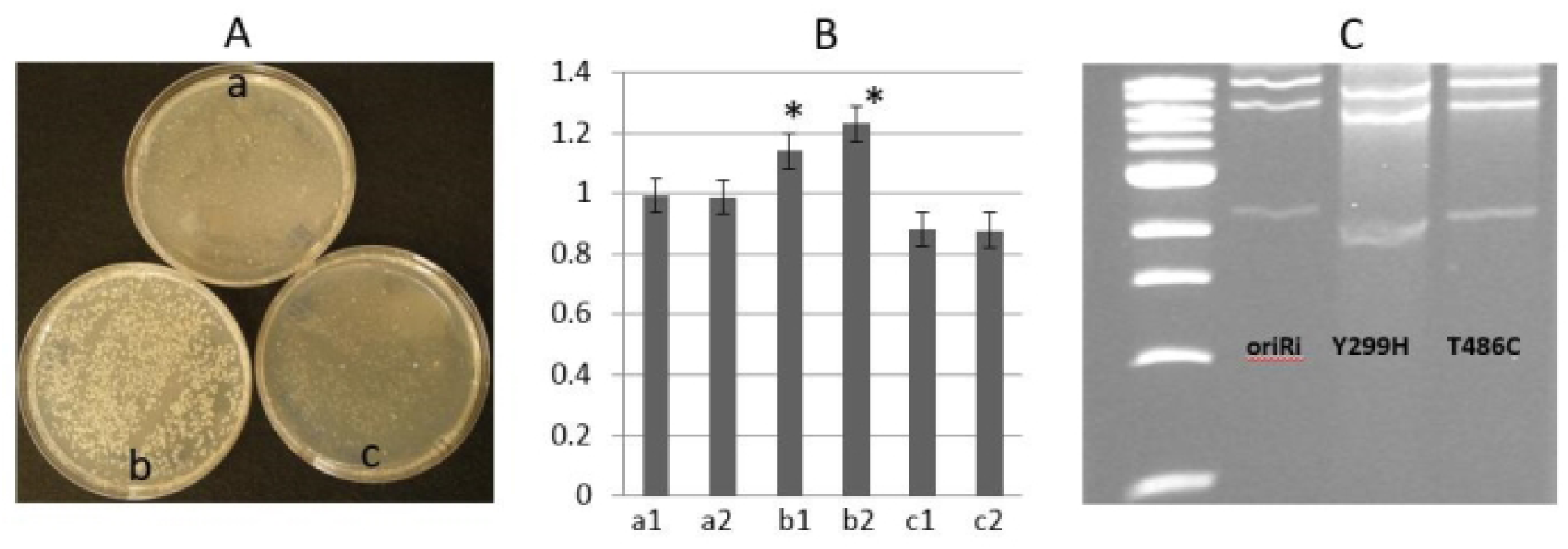
Comparison of oriRi binary plasmid growth and plasmid yield. Panel A) Colony size on solidified medium: a) pMON83937, WT-RepB; b) pMON83937-RepB^Y299H^; c) pMON8393-repB^T486C^ silent mutation; Panel B) Growth rate in liquid medium: a1, a2; two colonies from pMON83937, b1, b2; two colonies from pMON83937-RepB^Y299H^, c1, c2, two colonies from pMON8393-repB. Panel C) agarose gel showing plasmid yields from wild-type, RepB^Y299H^ and *repB*^T486C^ correspondingly. Error bars are shown in the diagram. *significant at P=0.01.

To check for increase in copy number of the plasmids with the mutations, cultures were grown to A_660_=1.0 and equal volumes were prepared for plasmid isolation. DNA from the preps was digested with *AflII* to give a distinct three band pattern (7.3 kb, 4.9 kb, 1.7 kb) on agarose gels. The gel picture clearly shows higher DNA concentration of each band on the gel from the RepB^Y299H^ cultures compared to the control (Figure 3C). DNA concentrations were not taken into consideration due to contamination with genomic DNA as observed on the gel. These observations indicate an increase in plasmid copy number resulting from the Y299H mutation.

To determine plasmid copy number in *Agrobacterium*, a DNA-based Taqman assay was employed using the single copy *Agrobacterium* chromosomal gene *lipA* as an internal copy number control. Atu0972 (lipA) was chosen as the internal control for comparison to the *gusA* sequence in the binary vectors because *lipA* copy number does not vary upon acetosyringone treatment [25]. A third *repB* double mutation plasmid, pMON83937 RepB^Y299H^ + *repB*^T486C^, was produced for copy number determination. As shown in Figure 4, the plasmid copy numbers varied between 1 (pMON83937, native oriRi) and 12 (pMON67438, multi-copy oriV binary vector) per cell. The oriRi RepB^Y299H^ mutant and the RepB^Y299H^ and *repB*^T486C^ double mutants were estimated to contain 9-10 copies/cell (Figures 4D and F), whereas the oriRi *repB*^T486C^ silent mutation alone showed approximately 1 copy/cell (Figure 4E).

**Figure 4:**
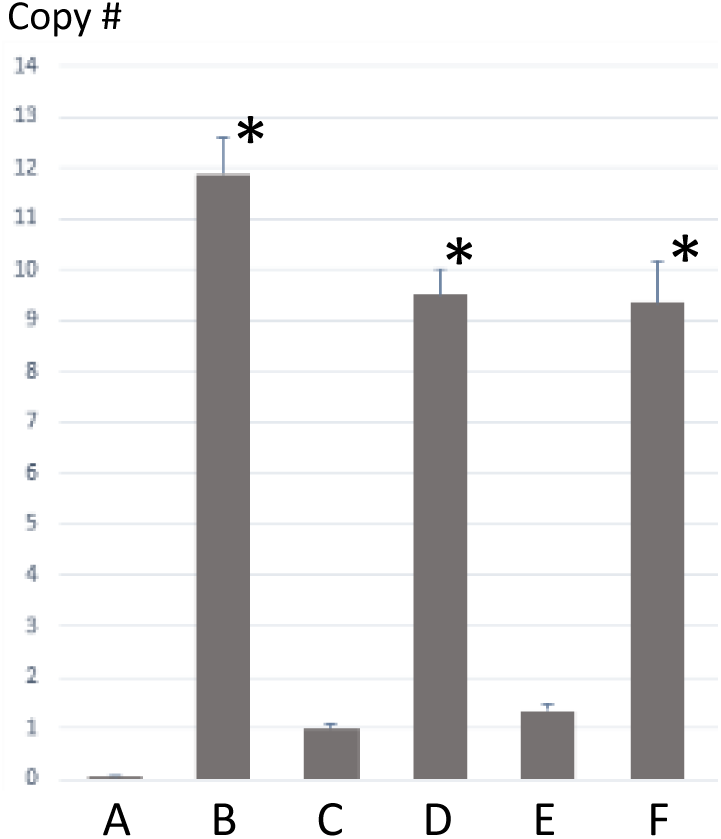
*Agrobacterium* plasmid copy number determination by ddPCR. A) *A. tumefaciens* no binary plasmid control; B) pMON67438 oriV replicon; C) pMON83937 oriRi native *repB*; D) pMON83937 RepB^Y299H^; E) pMON83937 repB^T486C^; F) pMON83937 RepB^Y299H^ + repB^T486C^ double mutations. Error bars are shown in the diagram. *significant at P = 0.01.

The increased copy number of the oriRi with the RepB^Y299H^ mutation in *Agrobacterium* may result from decreased RepB^Y299H^ binding activity to the inverted repeat (IR) in RepD [26]. The C-terminus of RepB from *Rhizobium leguminosarum* is responsible for RepB dimerization/oligomerization [19], and is conserved among the *repABC* family (Figure 2). However, whether the RepB^Y299H^ mutation in our oriRi binary vector affects dimerization has not been experimentally determined. It is less likely that the RepB^Y299H^ mutation reduces RepB interaction with RepA because the N-terminus of RepB is required for RepA interaction [19]. The *repABC* promoter 4 (P4) in *Agrobacterium* is repressed by RepA, whose activity is strongly enhanced by RepB [27]. The RepB^Y299H^ mutation may also reduce cooperative binding to *repABC* promoters, which further affects plasmid partitioning due to reduced oligomerization. The stability of the *repABC* replicon with the RepB^Y299H^ mutation was not determined because our plant transformation experiments were carried out with antibiotic selection, as was the case in previous experiments by Oltmanns et al. [14]. It is possible that the mutation affects dimerization/oligomerization, which partially blocks plasmid partitioning and leads to the accumulation of higher plasmid copy numbers in cells. Plasmid-free *Agrobacterium* cells may be avoided or minimized because the *Agrobacterium* used for plant transformation was grown using antibiotic selection.

The difference in plasmid copy number between Oltmanns et al. [14] and this study may result from the assay method, *Agrobacterium* strain, or growth time difference. Plasmid copy numbers are determined primarily by their replication origins, which show significant cell-to-cell variation within every population [28]. Oltmanns et al. [14] reported 15-20 copies/cell with pTF::Ri carrying RepB^Y299H^ and repB^T486C^ mutations in *A. tumefaciens* EHA105 using a DNA blot assay, whereas in this study we estimated that both the single and the double mutations in a nopaline-type ABI strain have 9-10 copies/cell, using a ddPCR assay. The DNA blot method likely has less resolution. The control vector RK2 oriV vector was estimated to be 7-10 copies/cell by DNA blot [14] [14] and 12 copy/cell by ddPCR in this study. In previous work we estimated the control RK2 oriV vector in ABI strain is 4-8 times relative to the native oriRi vector signals by DNA blot analysis [9]. The RK2 oriV copy number determined by ddPCR is close to the RK2 oriV-containing plasmid pSoup which was determined by a real time PCR method [10].

### The wild-type oriRi outperforms the RepB^Y299H^ mutated oriRi for generating single copy, backbone-free transgenic events in canola transformation

Due to the contradictory observations in single copy, backbone-free plant production between the two versions of oriRi binary vectors [9,14], we speculated that the higher copy number of pTF::Ri [14] with the RepB^Y299H^ mutation abolished the advantage of the native oriRi binary vector for production of single copy transgenic plants. To test this hypothesis, we compared the wild-type oriRi binary vector (pMON83937) and the RepB^Y299H^ mutant (pMON138207), as well as the multi-copy oriV binary vector (pMON67438), in canola transformation. In total, 1262 transgenic canola events were generated for all three constructs, and all three constructs showed statistically equivalent transformation frequencies, which is defined as the number of independent transformed plants produced per number of explants used (Figure 5). Both the single copy and the single copy, backbone-free transgenic plant frequencies were equivalent between the multi-copy oriV binary vector and the oriRi vector with RepB^Y299H^ high copy mutant, whereas the native single copy oriRi vector showed a significant improvement in the frequency of plants with a single copy of the transgene and absence of vector backbone sequences. The frequency of plants containing vector backbone sequences was decreased three-fold using the native oriRi vector compared to the oriV and the high copy oriRi mutant vectors.

**Figure 5:**
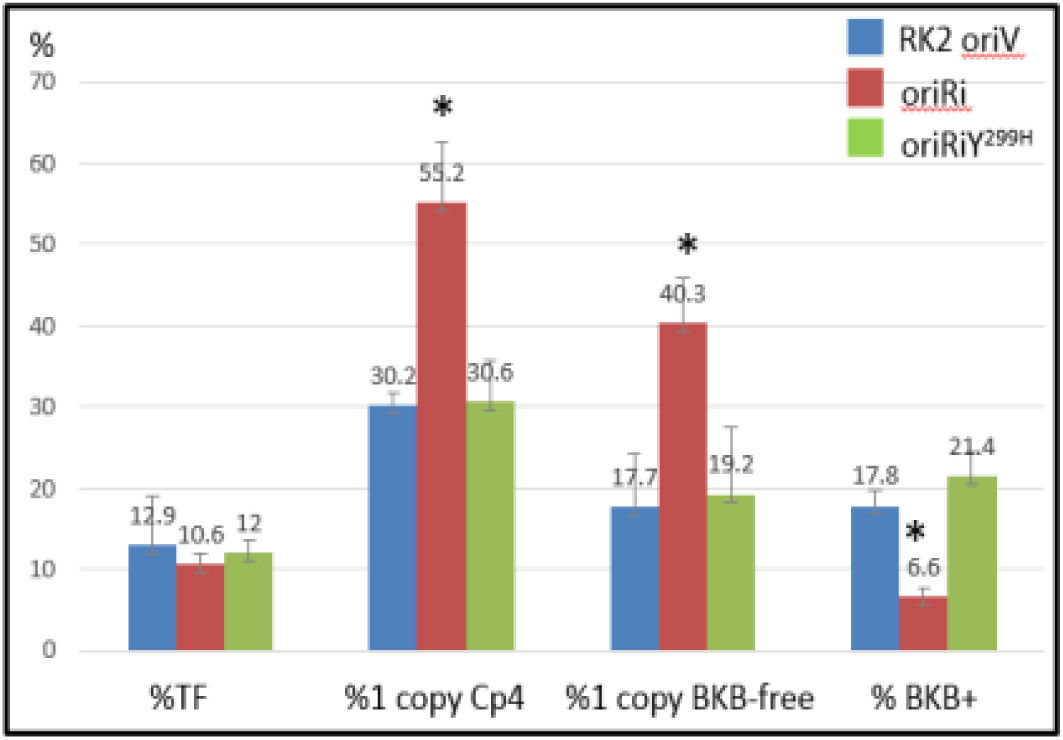
Effect of the oriRi with RepB^Y299H^ mutation on quality transgenic canola plant production. Total oriV (pMON67438) events n=421; total oriRi (pMON83937) events, n=379; total oriRi RepB^Y299H^ (pMON138207) events n=462. Error bars are shown in the diagram. *significant at P = 0.01. TF: transformation frequency (transgenic events/explant number); 1 copy CP4: single copy CP4 positive frequency; 1 copy CP4 BKB-free: single copy CP4 positive, backbone-free transgenic plant frequency; BKB+: backbone positive transgenic plant frequency.

This canola transformation result is consistent with the earlier observation that the high copy pTF::Ri resulted in a similar frequency in single copy, backbone-free plants to the multi-copy RK2 oriV binary vectors [14]. Our results support the hypothesis that T-DNA border processing involves a stoichiometric reaction between the endonuclease VirD2 and the border elements, and an excess of borders may titrate-out VirD2, resulting in incomplete processing [9,29]. This hypothesis is also supported by experiments in which a T-DNA was integrated as a single copy into an *Agrobacterium* chromosome, which drastically decreased *Agrobacterium* chromosome integration frequency in transgenic plants compared to other multi-copy binary vectors [14].

## Conclusion

Our data confirmed that the RepB C-terminus is critical for *repABC* replicon partitioning [19]. A single nucleotide mutation at RepB^Y299H^ in the ori pRi derived plant transformation binary vectors is responsible for an increase in plasmid copy number in *Agrobacterium*, and a concomitant decrease in relative frequency of single copy, backbone-free transgenic plants produced. This result also cautions biotechnological researchers that sequence confirmation is required for critical experiments and applications, since a single nucleotide mutation could lead to a very different conclusion.

## Author contribution statement

LG, SBG and XY conceived and coordinated the research. ZV made all plasmid mutagenesis. SJ created the original multi-copy ori pRi plasmid. SR and MLR conducted canola transformation. EN performed plasmid copy number estimation. ZV, SR, EN and XY analyzed the data. XY and ZV drafted the manuscript. All authors read and approved the manuscript.

## Acknowledgment

We thank our colleagues at the Monsanto Genome Sequencing Center for sequencing the pRi::Ti plasmid and all *repB* mutants, Drs. David Somers and Doug Boyes for supporting this research, and Dr. Charles Armstrong for critical review of the manuscript.

## Compliance with ethical standards

